# FhlA is a Formate Binding Protein

**DOI:** 10.1101/2024.07.24.604796

**Authors:** Abrar Abdullah Al Fardan, Benjamin James Koestler

## Abstract

*Escherichia coli* uses glycolysis and mixed acid fermentation and produces formate as by product. One system *E. coli* uses for formate oxidation is formate hydrogen lyase complex (FHL). The expression of the FHL complex is dependent on formate and regulated by the transcriptional regulator FhlA. The structure of FhlA is composed of three domains. The N-terminal domain is putatively responsible for formate binding and FhlA oligomerization as a tetramer, the central portion of FhlA contains a AAA+ domain that hydrolyzes ATP, and the C-terminal domain binds DNA. Formate enhances FhlA-mediated expression of FHL; however, FhlA direct interaction with formate has never been demonstrated. Formate-protein interactions are challenging to assess, due to the small and ubiquitous nature of the molecule. Here, we have developed three techniques to assess formate-protein interaction. We use these techniques to confirm that FhlA binds formate in the N-terminal domain *in vitro*, and that this interaction is partially dependent on residues E183 and E363, consistent with previous reports. This study is a proof of concept that these techniques can be used to assess other formate-protein interactions.

## Introduction

The bacterium *Escherichia coli* exhibits remarkable metabolic plasticity. During anaerobic growth and when there are no alternative electron acceptors, *E. coli* uses glycolysis and mixed acid fermentation, typically consuming C6 carbon like glucose and producing energy, succinate, acetate, lactate, ethanol, and formate [1]. This formate can be used as an electron donor for the electron transport chain [2,3], and this requires transport across the inner-membrane by the designated formate transporters FocA and FocB [4–8]. *E. coli* encodes three different formate oxidation systems: formate dehydrogenase N (encoded by the *fdn* operon)[9], formate dehydrogenase O (encoded by the *fdo* operon)[10,11], and formate hydrogen lyase (FHL, encoded by *fdhF* gene and the *hyc* operon)[12]. The two formate dehydrogenases (FDH-N and FDH-O) are responsible for the oxidization of periplasmic formate, and couple formate oxidation to nitrate reduction [13,14]. FDH-N is expressed during anaerobic growth and induced by nitrate, while FDH-O is active in the presence of oxygen or nitrate [13,14].

In contrast to the formate dehydrogenases, FHL is responsible for the disproportionation of cytoplasmic formate into CO_2_ and H_2_ [15,16]. Proteins of the FHL complex are encoded by the *hyc, hyp,* and *hyf* operons, the *fdhf* gene, and the *hyd* locus [12,17]. Expression of these genes is induced by formate under anaerobic conditions, dependent on the NtrC family transcriptional regulator FhlA [18]. FhlA binds the upstream region of the *fdhF* and the *hyc* operons [19,20]. The structure of FhlA is composed of three domains [21,22](Fig. 1). The FhlA N-terminal domain consists of amino acids 1-381 and is believed to facilitate homooligimerization as a tetramer [22]. FhlA encodes a AAA+ domain in the middle of the protein (amino acids 388-617) that hydrolyzes ATP and interacts with RNA polymerase and σ^54^ to initiate transcription [22,23]. And at the C-terminus of the protein (amino acids 618-692) is a helix-turn-helix domain that binds the one side of the DNA upstream of FHL genes [22]. For a comprehensive description of formate metabolism and its underlying genetics, refer to this review [24].

**Figure 1.**
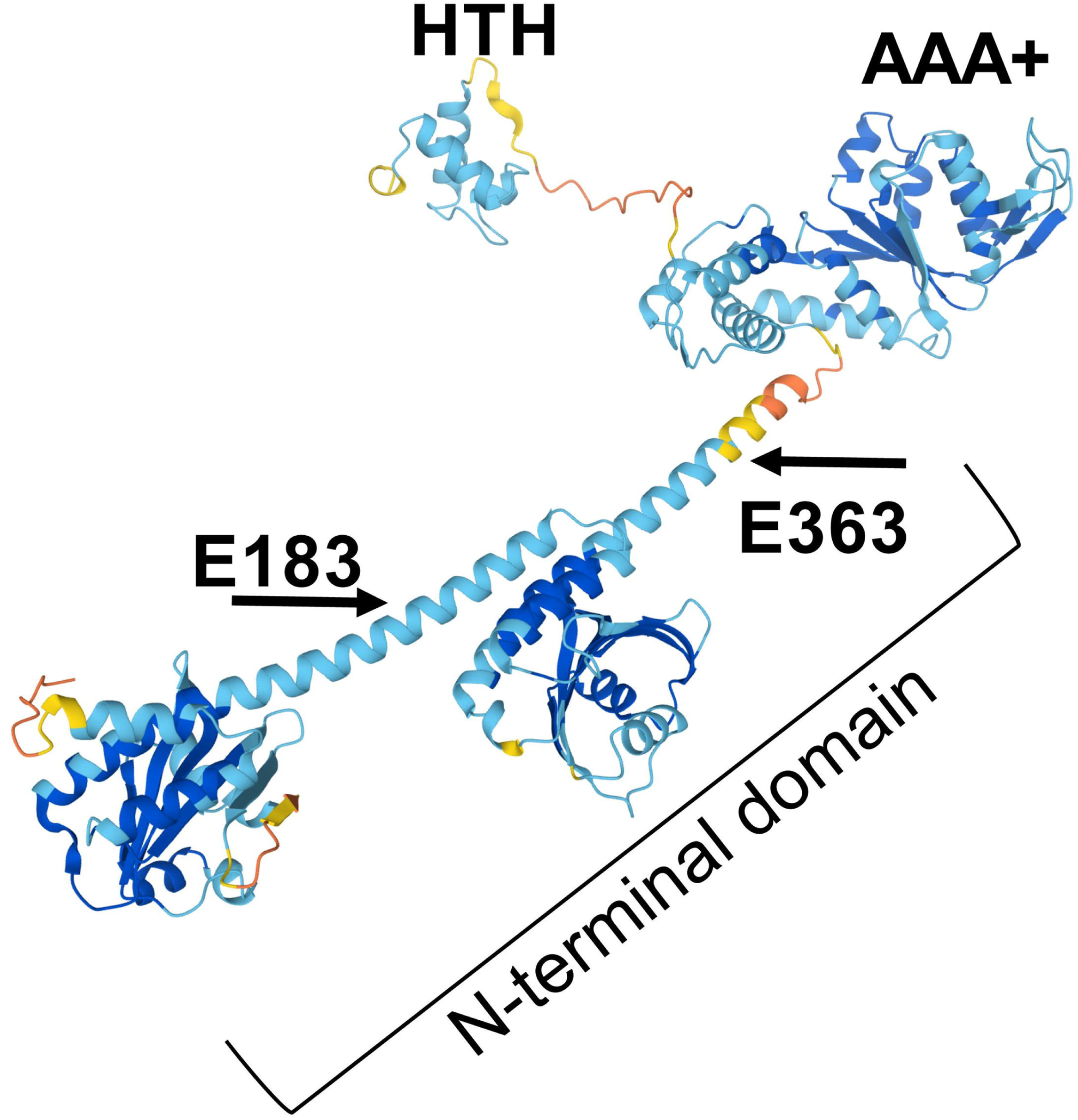
FhlA is a DNA-binding transcriptional regulator, composed of three domains. The N-terminal domain facilitates tetramerization and is the putative formate-binding domain; the location of E183 and E363 residues are highlighted. The central portion of the protein is a AAA+ domain, which hydrolyzes ATP. And the C-terminal portion is a helix-turn-helix domain, which interacts with DNA. The predicted structure of FhlA was determined by Alphafold; color indicates pLDDT confidence, with darker blue indicating high confidence structural prediction, while yellow and orange indicate low confidence [25,26].

Formate increases FhlA-dependent expression of *fdhF* and *hyc* [22,23,25,26], but how formate interacts with FhlA is unknown. There are several lines of evidence that support formate binding occurs in the FhlA N-terminal domain. Although formate enhances ATP hydrolysis of the AAA+ domain [23], deletion of the FhlA N-terminal domain causes formate-independent and constitutive transcription of FHL genes [21]. Furthermore, an *fhlA* E183K mutation (located in the N-terminal domain) constitutively activates *hyc* transcription regardless of formate, whereas a E363K mutation (also in the N-terminal domain) exhibits semi-constitutive *hyc* expression in response to formate [27]. While these genetic studies show that formate enhances FhlA-mediated expression of FHL, there has been no direct demonstration that FhlA directly interacts with formate. As formate is a low molecular weight molecule containing a single carbon, there are few analytical methods to measure formate, and no characterized protein domains are known to interact with formate. Here, we developed three techniques to assess formate-protein interactions, using ^14^C-labelled formate for sensitive detection. We use these techniques to confirm that FhlA binds formate in the N-terminal domain *in vitro*, and that this interaction is partially dependent on residues E183 and E363, confirming previous reports. These techniques can be applied to study other formate binding proteins.

## Materials and methods

### Media and culture conditions Strains and plasmids

Bacterial strains, plasmids, and primers used in this study are listed in Table 1. *E. coli* BL21 (DE3) with a Δ*fhlA*::Kan mutation was used as a host for plasmid cloning, which was generated using P1 transduction [28]. *E. coli* MG1655 full-length FhlA and FhlA-NT expression plasmids were generated by PCR and restriction cloning into pET22b using HindIII and NdeI sites [29]. Plasmid sequence was confirmed by sequencing (Azenta). The E183K and E363K mutations were generated by site-directed mutagenesis, using Phusion polymerase and primers containing the missense mutations. *E. coli* strains were routinely cultured at 37°C on Luria-Bertani broth (LB). *E. coli* BL21 carrying plasmids encoding FhlA-NT, FhlA-NT E183K, and FhlA-NT E363K mutations were always grown with 25 μg/ml ampicillin; for purification of full-length FhlA, the chaperone plasmid pG-KJE8 was included [30,31], and these cultures were supplemented with additional 20 μg/ml chloramphenicol, 0.5 mg/ml L-arabinose, and 1 ng/ml tetracycline.

**Table 1:** Strains and plasmids used in this study.

| Strain or plasmid | Description |
| --- | --- |
| *Escherichia coli* strains |  |
| BL21 $\Delta fhIA::Kan$ | |

| Plasmids |  |
| --- | --- |
| **pET22b** | Parental plasmid for *fhlA* expression strains |
| **pG-KJE8** | Chaperone for protein purification [31] |
| **pfhlA** | Full length *fhlA* in pET22b |
| **pfhlA-E183K** | Full length *fhlA* with E183K in pET22b |
| **pfhlA-E183K** | Full length *fhlA* with E363K in pET22b |
| **pfhlA-NT** | *fhlA* N-terminal portion (1-381) in pET22b |
| **pfhlA-NT-E183K** | *fhlA* N-terminal portion (1-381) with E183K in pET22b |
| **pfhlA-NT-E363K** | *fhlA* N-terminal portion (1-381) with E363K in pET22b |

### Protein purification

Purification of FhlA derivatives was similar to Schlensog et al. (1994) with modifications [32]. Overnight cultures were subcultured in 1.5 L of protein expression media (12 g/L tryptone, 24 g/L yeast extract, 40 ml/L glycerol, 2.13 g/L K_2_HPO_4_, and 12.54 g/L KH_2_PO_4_ , pH 7.2)[33] and grown at 37°C with shaking (aerobic) to an optical density (OD_600_) of 0.6 to 0.8 (mid-log). Cultures were then induced with 100 μM IPTG and grown overnight at 20°C. Cultures were centrifuged to pellet, and resuspended in 35 ml of buffer A (20 mM imidazole, 50 mM KH_2_PO_4_, 300 mM NaCl, 10% glycerol, 2M KCl at pH 8). EDTA-free protease inhibitor cocktail and nuclease were added, then cells were lysed on ice by sonication. Cell debris was removed by centrifugation at 19000 x g for 15 minutes, and then the supernatant was run through nickel affinity column. Buffer exchange was performed using BioRad 10DG desalting columns (Cat. 7322010) according to the manufacturer’s instructions.

The protein content of these preparations was analyzed by both Coomassie stain and western blot. Samples were mixed with loading buffer (375 mM Tris-HCl pH6.8, 9%SDS, 50% glycerol, 9% β-mercaptoethanol, and 0.03% bromophenol blue) and boiled for 10 min and then electrophoresed in duplicate SDS-PAGE gels (10% acrylamide/bis, 37.5:1). After electrophoresis, one gel was stained with Coomassie blue, and the protein from the other gel was transferred to a 0.45-mm-pore-size nitrocellulose membrane (Hybond-ECL; GE Healthcare) and incubated with mouse α-HIS antibody (Genscript A00186). Proteins were detected using HRP-conjugated goat α-mouse antibody (Abclonal AS003), and a Pierce ECL detection kit (Thermo-Fisher Scientific).

### Formate Pulldown Assay

Purified protein was bound to Ni-NTA Magnetic Agarose Beads (ThermoFisher 78605), according to the manufacturer’s instructions with modification. 500 μL of protein sample was prepared by mixing 125 μL of equilibration buffer (50 mM Na_3_PO_4_, 0.3 M NaCl, 10 mM Imidazole, 0.05% tween at pH 8), 125 μL wash buffer (50 mM Na_3_PO_4_, 0.3 M NaCl, 15 mM Imidazole, 0.05% tween at pH 8), and 250 μL of purified protein (32 μM final concentration). For the no-protein control, equimolar histidine was used. 50 μL of bead slurry equilibrated by washing twice with 400 μL of equilibration buffer using a magnetic stand, and then 250 μL of prepared sample or no-protein control was mixed with 6 μCi (0.5 μM) ^14^C-labeled formate and added to the beads. Tubes were vortexed and mixed on an end-over-end rotator for one hour at RT. The beads were washed twice with 250 μL of wash buffer, and then 125 μL of elution buffer (50 mM Na_3_PO_4_, 0.3 M NaCl, 0.3 mM Imidazole at pH 8) was added and mixed on an end-over-end rotator for 20 minutes. The beads were removed using a magnetic stand and the supernatant was transferred to scintillation cocktail (Fisher 19144) and quantified by scintillation (Tri-Carb 4910TR liquid scintillation counter).

### Equilibrium dialysis assay

FhlA-NT, E183K, and E363K derivatives (32 μM each) were mixed with 6 μCi (0.5 μM) ^14^C-labeled formate in equilibration buffer (10 mM Tris-pH 8, 100 mM NaCl and 5 mM MgCl_2_) at a final volume of 50 μL. The mixture was placed in a two-chamber equilibrium dialysis system with a regenerated cellulose membrane (5000 Da MWCO, Harvard Bioscience 74-2200), containing 50 μL equilibration buffer in the opposite chamber. Samples were collected over time from the no-protein chamber and suspended in ScintiVerse™ BD Cocktail (Fisher SX18-4), and quantified using a scintillation counter (Tri-Carb 4910TR).

### DRaCALA assay

^14^C-formate DRaCALA was performed similarly to c-di-GMP DRaCALA, with some modification [34]. ^14^C-labeled formate was mixed with DRaCALA binding buffer (10 mM Tris-pH 8, 100 mM NaCl and 5 mM MgCl_2_) to a final concentration of 60 μCi (1 μM); 20μL of this formate mixture was added to 50 μL of protein (32 μM) and incubated on ice for 10 minutes. 10 μL protein-formate mixture was spotted on a 0.2 μm dry nitrocellulose membrane, in triplicate. The membrane was dried completely then used to expose a phosphor screen, and visualized with a phosphorimager (Personal Molecular Imager (PMI) System, Bio-Rad 170-9400). The data were quantified using ImageJ. The fraction bound between all the samples was calculated as follows where I= intensity and A=area [34]:

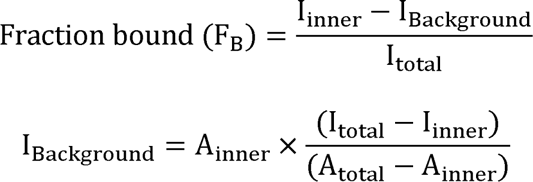

### Statistical Analysis

Statistical analysis was performed using Graphpad Prism.

## Results

### Formate binds to the FhlA N-terminal domain

To determine if FhlA directly interacts with formate, we examined *E. coli* K-12 MG1655 FhlA (protein ID: NP_417211.1) *in vitro*. We purified the N-terminal portion of FhlA (AA 1-381) to determine if formate binds this portion of the protein, hereon referred to as FhlA-NT (Fig. 1). The His-tagged FhlA-NT was purified using immobilized metal-ion affinity chromatography, and we confirmed FhlA-NT by SDS-PAGE followed by Coomassie staining and western blotting (Fig. 2). Purification of each NT fragment was consistent, with minor differences in efficiency reflected in band intensity. First, we performed a ^14^C-formate pulldown assay using the purified FhlA-NT. We used magnetic nickel nitrilotriacetic acid agarose beads (Ni-NTA), where His-tagged proteins were bound to Ni-NTA magnetic beads, mixed with ^14^C-labeled formate, and bound ^14^C - labeled formate was quantified by liquid scintillation. Of note, because formate is a ubiquitous biomolecule, and because there are no previous studies that have defined a protein that does not interact with formate, we opted to use a no-protein treatment as a negative control. Scintillation counts for FhlA-NT were significantly higher when compared to the no-protein control (Fig. 3A). This indicates ^14^C-formate binds FhlA between residues 1-381. In subsequent repetitions of this assay, scintillation counts from the FhlA-NT treatment were consistently higher than the no-protein control, but the extent of this increase varied widely (Fig. 3B); therefore, we sought to develop other methods to assess formate-protein interactions.

**Figure 2.**
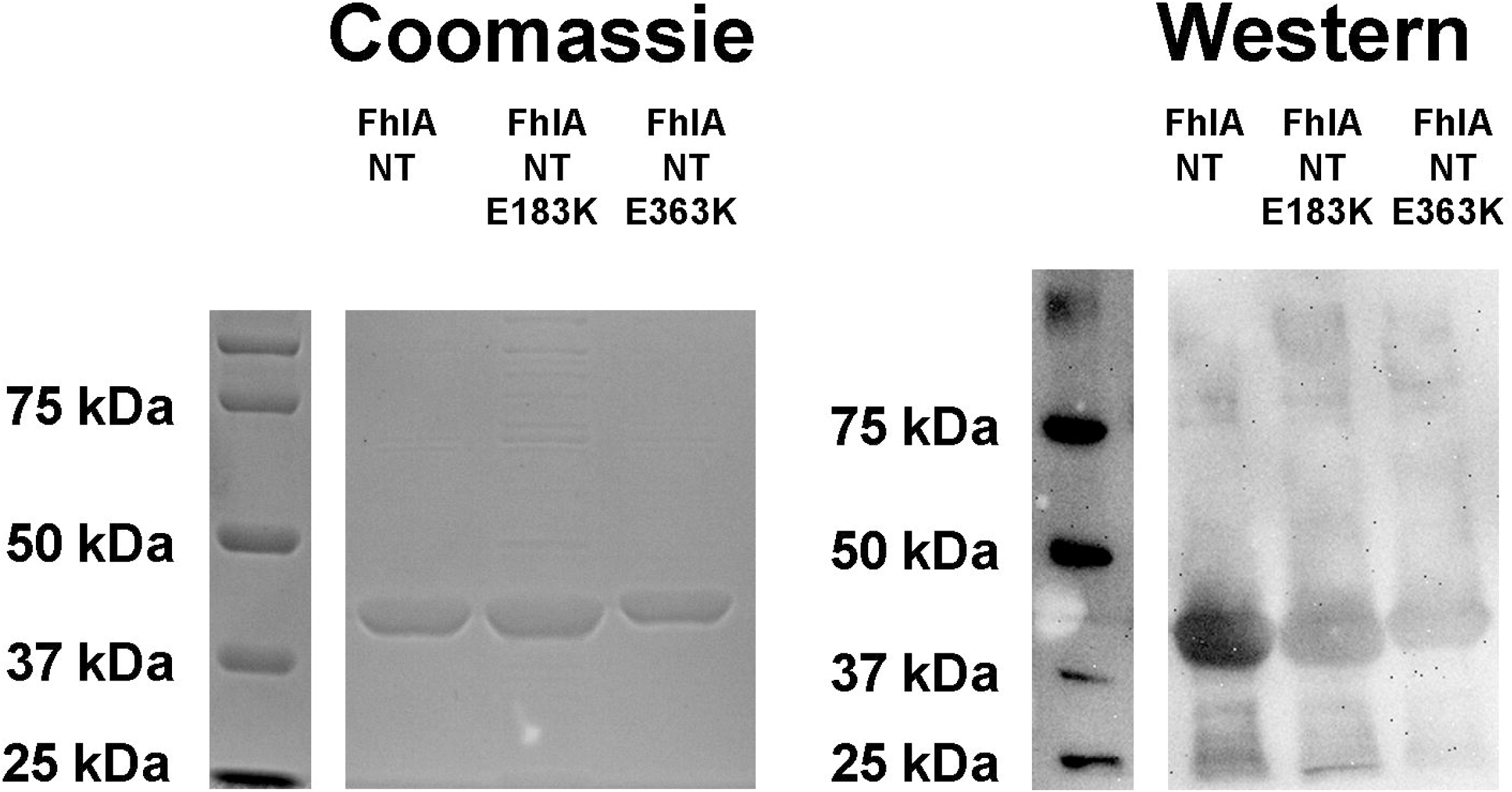
Purification of FhlA-NT was confirmed by Coomassie staining (left) and western blotting (right). Proteins were run on a 10% SDS-PAGE gel. Corresponding ladders are shown on the left. We observed a band at 43 kDA, consistent with the size of FhlA-NT. The E183K and E363K derivatives also showed similar sized bands. Antibodies used in the western blot are targeting the HIS tag. Samples were not normalized prior to loading in each gel.

**Figure 3:**
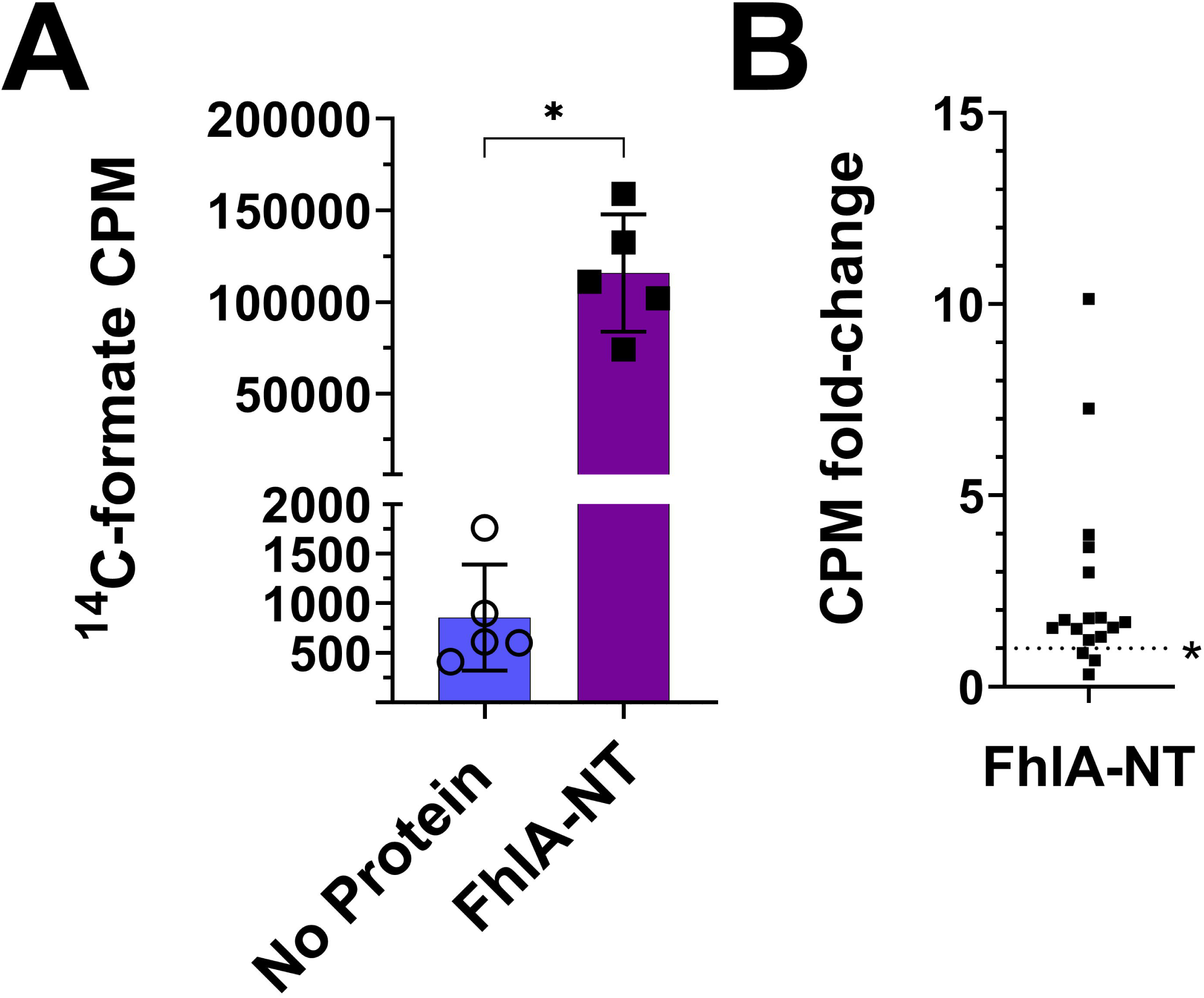
FhlA-NT bound to Ni-NTA beads retains ^14^C-labeled formate. (A) The counts per minute for FhlA-NT were significantly higher compared to a no protein control. Asterisks represent significant differences between treatments, as determined by a T- test (n = 5, p<0.05). Symbols indicate individual replicates; error bars indicate standard deviation. (B) In subsequent replications, repeated on 7 different days, we observed a significant increase in CPM of FhlA-NT compared to the no-protein control (single sample T-test compared to 1, n=17, p < 0.05), but the fold-change varied widely.

### E183K mutation reduces FhlA formate binding

Because *fhlA* E183K and E363K mutations impact the ability of FhlA to regulate FHL expression in response to formate [27], we generated E183K and E363K mutant derivatives of FhlA-NT. To test formate interaction, we performed equilibrium dialysis [35,36], where a mixture of ^14^C-formate and FhlA-NT is added to a two-chamber system separated by size exclusion membrane. Samples were collected every hour for 10 hours from the no-protein chamber and quantified by liquid scintillation to track the diffusion of formate away from protein. We observed that FhlA-NT caused significantly lower ^14^C-formate diffusion across the membrane when compared to no-protein (Fig. 4). The FhlA-NT E363K mutant had similar diffusion pattern across the membrane to FhlA-NT (Fig. 4). In contrast, the diffusion of ^14^C-formate in FhlA-NT E183K samples were comparable to the no-protein control, indicating that E183 contributes to FhlA-formate interaction.

**Figure 4.**
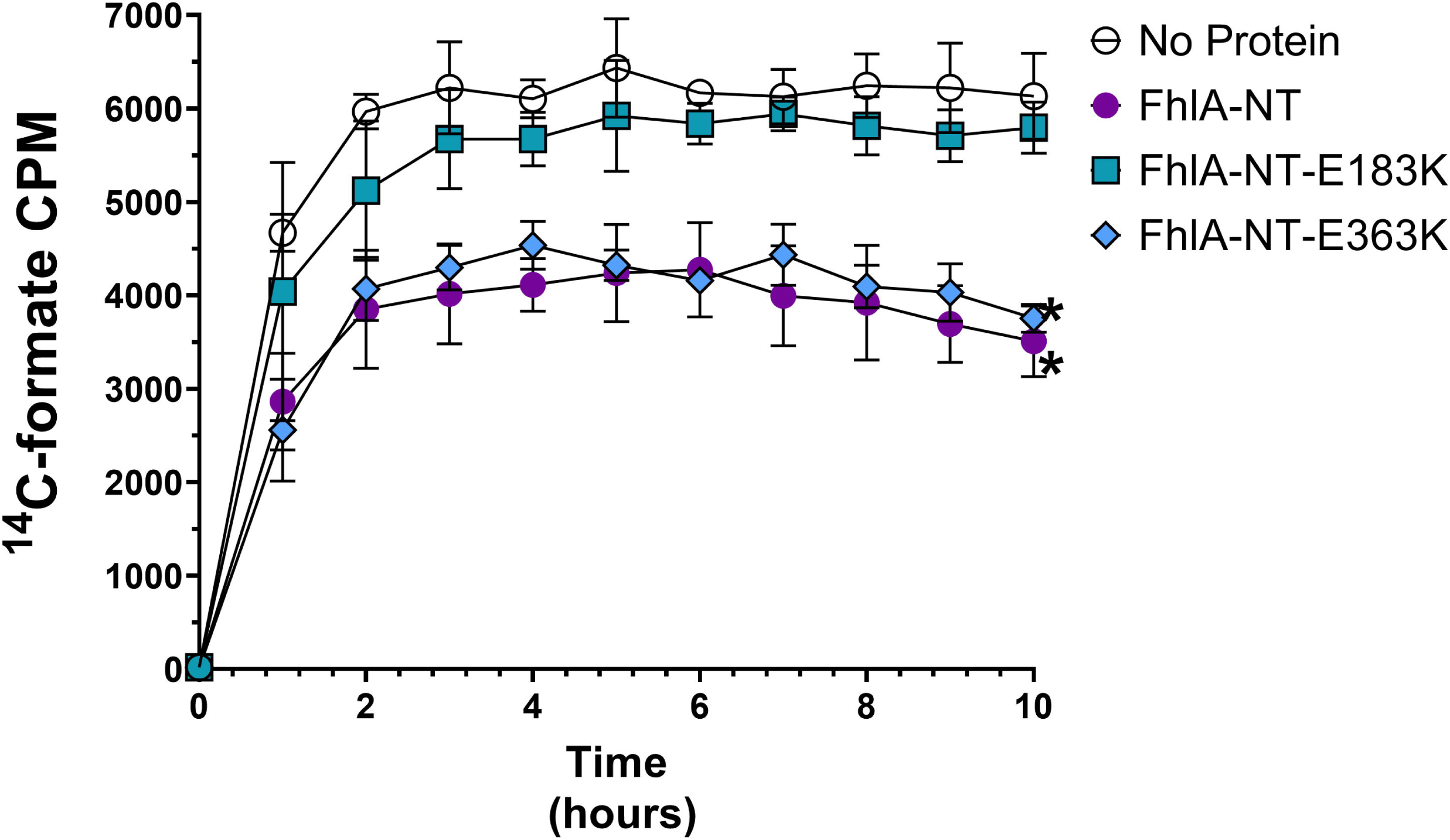
FhlA-NT-E183K is unable to retain formate when dialyzed. Proteins were mixed with^14^C-labeled formate and equilibrium dialysis was performed. Samples were collected from the no-protein chamber. FhlA-NT and FhlA-NT-E363K caused a slower diffusion across the membrane than the FhlA-NT-E183K mutant protein or the no protein control. Asterisks represent significant differences between proteins, as determined by two-way ANOVA (n=3, p<0.05). Error bars indicate standard deviation.

### A FhlA E363K mutation reduces formate binding in a DRaCALA assay

As an alternative approach to assess FhlA-NT interaction with formate, we used differential radial capillary action of ligand assay (DRaCALA)[34]. DRaCALA is based on the mobility difference of free ligand and protein-bound to the radiolabeled ligand, and has been used to study other protein-ligand interactions including c-di-GMP, c-di-AMP, cAMP, and (p)ppGpp [34,37–41]. We performed DRaCALA using ^14^C-formate and compared the FhlA-NT to the E183K and E363K mutant derivatives. We found that the relative ^14^C-formate fraction bound of the FhlA-NT was significantly higher than the no protein control, consistent with FhlA interaction with formate. Likewise, the FhlA-NT fraction bound was also significantly higher than FhlA-NT E183K and E363K derivatives, indicating both FhlA-E183K and FhlA-E363K proteins are deficient in formate binding. However, the FhlA-NT E363K protein had lower fraction bound compared to the E183K, suggesting that this mutation was more detrimental than E183K for formate-binding. The E183K mutation also had higher formate binding than the no-protein control, which is a different pattern than we observed using equilibrium dialysis. DRaCALA with full-length purified FhlA, and full-length E183K and E363K derivatives produced similar pattern to that of the FhlA-NT proteins (Fig. 4).

## Discussion

There is significant interest in bacterial formate metabolism, as it can be harnessed to produce hydrogen in a renewable manner [19,42–44], or used to capture atmospheric CO_2_ [45]. Formate is also a common biomolecule, and contributes to many notable biological processes including cancer metabolism and bacterial pathogenesis [46–50]. However, formate is particularly challenging to study due to its small and ubiquitous nature, and there is a lack of methodologies to effectively measure this molecule. For example, others have used indirect methods to characterize formate transport and metabolism [8]. One of the few known formate-binding proteins is FhlA, but formate interaction has never been directly demonstrated. FhlA is maximally active when formate is present in the low mM range [18,21,23,27,42,51]. We demonstrate here that formate directly binds to the N-terminal domain of FhlA, and this is partially dependent on E183 and E363. This is consistent with previous studies that show that formate promotes FhlA-mediated expression of *fdhF* and the *hyc* operon [21–23,25–27,51], essential for FHL function [52]. The FhlA N-terminal domain likely facilitates oligomerization of a homeric tetramer [42]. Our current hypothesis is that formate interacts with the N-terminal domain to alter the quaternary structure of FhlA and subsequently activate FhlA-dependent transcription; further experimentation is required to test this hypothesis.

Here, we present three new assays to assess formate-protein interactions. The formate-pulldown assay directly measures ^14^C-formate by scintillation from bound protein, allowing for higher sensitivity and lower input protein [53], but in the case of ^14^C-formate this assay was less consistent. Equilibrium dialysis allowed us to observe protein-formate interactions when protein is freely solubilized. The DRaCALA assay also showed binding between FhlA and formate; this assay is scalable as a high-throughput assay, but requires a higher amount of ligand for visualization by autoradiography [34]. Interestingly, the FhlA-NT E363K mutant showed formate binding in our equilibrium dialysis assay, but not in our DRaCALA assay (Fig. 4, 5). The most significant difference between these techniques is that protein is solubilized during equilibrium dialysis, whereas protein is immobilized during DRaCALA; we speculate that this impacts formate-binding by altering protein oligomerization. One limitation of this study is the lack of a functional assay for the FhlA-NT, meaning we cannot rule out that changes in protein conformation account for these differences. Another limitation was our lack of a protein negative control; however, our study provides evidence that FhlA and mutant derivatives can be used as controls for these assays in future studies. This study also highlights the value in using different methodologies to interrogate protein-ligand interactions.

**Figure 5.**
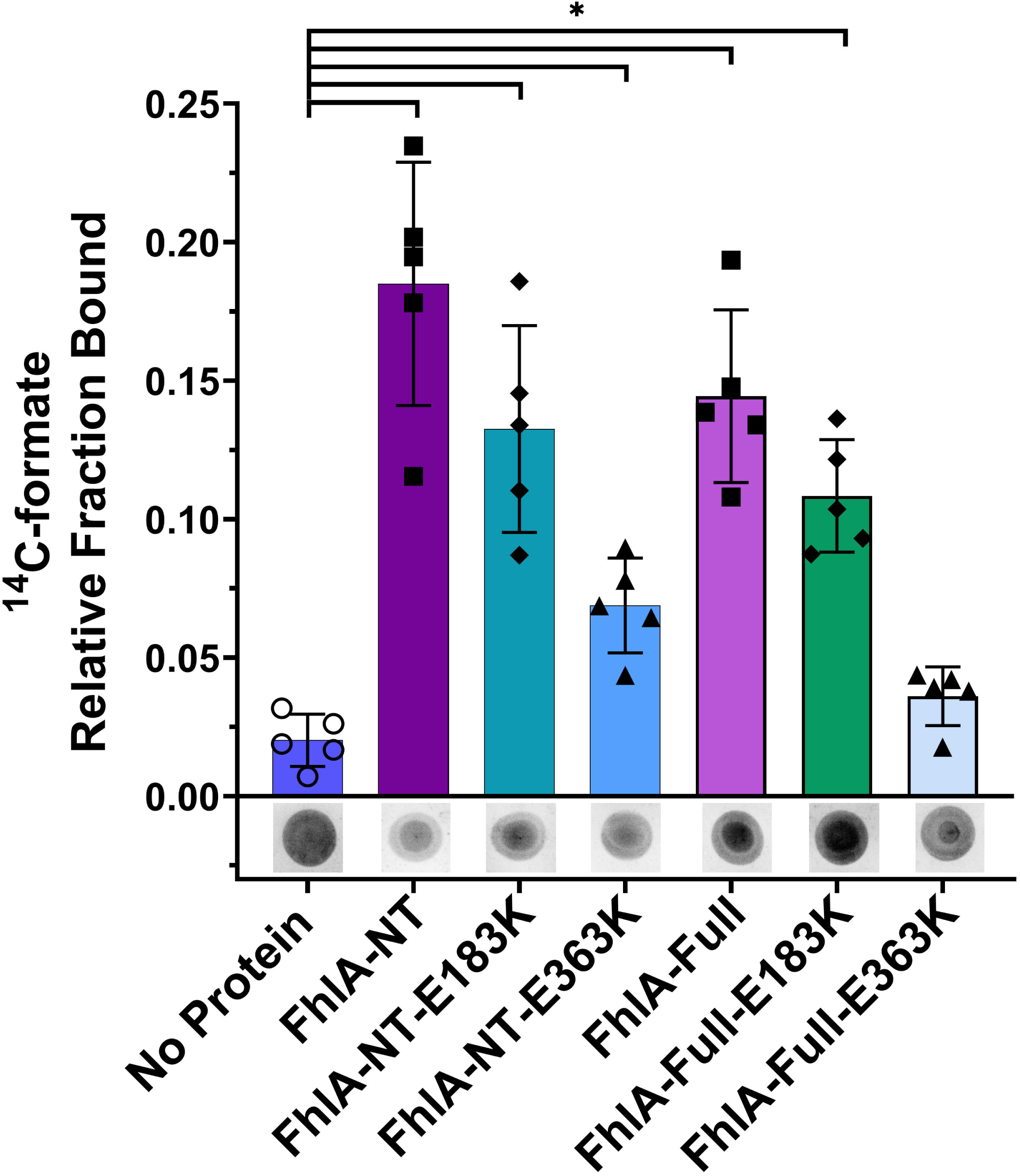
The FhlA-NT-E363K mutant derivative shows reduced DRaCALA formate binding. FhlA-NT, FhlA-NT-E183K, and FhlA-NT-E363K were mixed with^14^C-labeled formate and spotted on a nitrocellulose membrane; formate interaction was visualized by autoradiography. FhlA-NT showed the highest fraction bound, compared to a no protein control. The E183K and E363K mutant derivatives each showed lower formate binding, but still higher than the no-protein control. Asterisks represent significant differences between proteins compared to the no protein control, as analyzed by one-way ANOVA (n=5, p<0.05). DRaCALA spots are representative of the 5 replicates. Error bars indicate standard deviation

## Acknowledgements

We would like to thank Bobby Zhang and Dr. Ricky Stull for help purifying FhlA. This work was supported by the NIH R03 grant AI156432-01A1, and startup funds and a grant from the Faculty Research and Creative Activities Award, Western Michigan University.

## Data availability

We confirm that the data supporting the findings of this study are available within the article and its supplementary materials, and raw data are available from the corresponding author (BK) upon request.

